# Bayesian areal disaggregation regression to predict wildlife distribution and relative density with low-resolution data

**DOI:** 10.1101/2023.01.12.523783

**Authors:** Kilian J Murphy, Simone Ciuti, Tim Burkitt, Virginia Morera-Pujol

**Affiliations:** Laboratory of Wildlife Ecology and Behaviour, School of Biology and Environmental Science, University College Dublin, Ireland; DEER MANAGEMENT SOLUTIONS, Coolies, Muckross, Killarney, Co. Kerry V93R380

## Abstract

For species of conservation concern and human-wildlife conflict, it is imperative that spatial population data are available to design adaptive-management strategies and be prepared to meet challenges such as land use and climate change, disease outbreaks, and invasive species spread. This can be difficult, perhaps impossible, if spatially explicit wildlife data are not available. Low-resolution areal counts, however, are common in wildlife monitoring, i.e., number of animals reported for a region, usually corresponding to administrative subdivisions, e.g., region, province, county, departments, or cantons. Bayesian areal disaggregation regression is a solution to exploit areal counts and provide conservation biologists with high-resolution species distribution predictive models. This method originated in epidemiology but lacks experimentation in ecology. It provides a plethora of applications to change the way we collect and analyse data for wildlife populations. Based on high-resolution environmental rasters, the disaggregation method disaggregates the number of individuals observed in a region and distributes them at the pixel level (e.g., 5×5 km or finer resolution), therefore converting the low-resolution data into high-resolution distribution and indices of relative density. In our demonstrative study, we disaggregated areal count data from hunting bag returns to disentangle the changing distribution and population dynamics of three deer species (red, sika and fallow) in Ireland from 2000 to 2018. We show an application of Bayesian areal disaggregation regression method and document marked increases in relative population density and extensive range expansion for each of the three deer species across Ireland. We challenged our disaggregated model predictions by correlating them with independent deer surveys carried out in field sites and alternative deer distribution models built using presence-only and presence-absence data. Finding high correlation with both independent datasets, we highlighted the accurate ability of Bayesian areal disaggregation regression to capture fine scale spatial patterns of animal distribution. This study opens new scenarios for wildlife managers and conservation biologists to reliably use regional count data disregarded so far in species distribution modelling. Thus, representing a step forward in our ability to monitor wildlife population and meet challenges in our changing world.

**Open data statement:** Data used in the study has been publicly archived for reproducibility.

Data archive DOI: 10.6084/m9.figshare.21890505

## Introduction

The task of managing wildlife species and their habitats requires an empirical approach to confidently respond to scenarios such as land-use change, disease outbreaks, invasive species spread, among others. Decision-making frameworks ensure that managers in rapidly changing systems can act while acknowledging uncertainty and variability (e.g., strategic adaptive wildlife management, evidence-based conservation systems, see Gilson et al. 2019). Though these frameworks may differ in their philosophy and methodologies, they share the common theme of data-guided management (Gillson et al. 2019). Systematic wildlife monitoring schemes underpin decision making frameworks and are required to track wildlife populations’ response to shifting ecological conditions (Meyer et al. 2010). This can be difficult, perhaps impossible if spatially explicit wildlife point data are not available to inform managers of wildlife population traits in space and time (Lidenmayer and Likens, 2009). Low spatial resolution areal counts are commonly available for wildlife (e.g., number of animals harvested or simply observed for a region, province, county, or other administrative boundary), however these data lack spatial resolution, limiting their use in preference of higher resolution data such as observations from sophisticated tracking devices e.g., GPS, VHF (Thomas et al. 2013). Unfortunately, systematic monitoring schemes take time to become established, demand a significant amount of time and financial investment to remain viable, and require careful governance of the programme and curation of the data – facets that often-become barriers to the establishment and continuation of a successful systematic monitoring scheme (Lindenmayer and Likens, 2009; Murphy et al. 2022).

High-resolution data are generated by remote-sensing instruments (e.g., satellites), unmanned technology (e.g., drones, camera traps, acoustic monitors), smart devices (e.g. smartphones) and human observation (e.g. surveys, citizen science) by a diverse cast of citizen scientists, stakeholders, scientific and government agencies (Hampton et al. 2013). When these data are collected but not curated or analysed, they risk becoming “dark data”, stored indefinitely but not contributing to decision-making frameworks for management, thus wasting resources on collecting unexploited data and following uninformed management actions (Hampton et al. 2013). Data are difficult to use if not easily obtainable, lack metadata, are not stored in a digital format, or are indecipherable due to lack of interest in curating the data appropriately (Simpson et al. 2020). The challenges of data sharing often have a human dimension where nuance exists in cultural, societal, and professional norms that may limit different organisations (e.g., academia and government bodies) communicating common goals and working in tandem to achieve them (Michener and Jones, 2012; Murphy et al. 2022). The emergence of online data repositories (e.g. Movebank; Dryad), the adoption of standard data protocols and good data stewardship practices, paired with cultural change such as, increased informatics literacy, data sharing, and awareness of open science benefits has improved how we handle data, however, “dark-data” are still prevalent when historically collected data are stored in unknown silos (Michener and Jones, 2012; Hampton et al. 2013). Novel statistical methods offer an opportunity to revitalise dark data, for instance, low-resolution data, which may now be capable of informing wildlife management in the absence of high-resolution spatial data.

One such emerging method is Bayesian areal disaggregation regression. Although relatively unknown in ecological circles, it is an emerging method in statistical epidemiology, capable of providing “pixel-level” predictions (typically at 1×1km^2^ or 5×5km^2^) from data collated at the regional level. Such regional-level data are usually the number of animals observed or harvested within administrative borders, such as region, province, county, departments, or cantons. To obtain high-resolution predictions from aggregated areal data (i.e. wildlife observations by jurisdiction), disaggregation regression uses fine-scale covariate data and models the likelihood of the response data as the weighted sum of these processes in each aggregation unit (Arambepola et al. 2021). The use of high-resolution covariate information, together with predefined jurisdictional borders helps avoid common pitfalls related to the change of scale such as the “modifiable areal unit problem” (i.e., statistical bias when samples are used to represent information such as density in a given area, see Fotheringham and Wong, 1991). Simulation studies have shown promise but have raised questions about the appropriate usage of this method in real world scenarios (Arambepola et al. 2021). While not previously used in an applied ecological context it has been successfully applied to disease risk modelling for COVID-19 (Python et al. 2021) and malaria (Lucas et al. 2021). Bayesian areal disaggregation regression has the potential to be readily used in ecology due to the availability of high-resolution environmental data from remote sensing (Nagendra et al. 2013) and abundant low resolution areal wildlife count data (e.g., bird and mammals census or harvest statistics at the regional level).

Ireland provides an excellent case study to evaluate this methodology, Ireland hosts a mixture of native red deer (*Cervus elephus*), and non-native introduced species, sika (*Cervus nippon*), and fallow (*Dama dama*) deer and lacks a national systematic monitoring programme or management plan for any deer species. Hunting bag return data reported at the county level is the sole metric of deer population dynamics over time available for the last 20 years. There are many interested stakeholder groups (e.g., farmers, foresters, hunters, conservationists) that view these species as either a pest or a resource, and while management philosophies exist across this spectrum, the shared management goal of these stakeholder groups is the presence of healthy and sustainable ungulate populations. Since 2000, Ireland has experienced rapid and expansive land-use change, coupled with suspected deer territorial expansion and population growth (Carden et al. 2011) that has been implicated in the emergence of a myriad of human-deer coexistence issues e.g. crop and forest damage, disease transmission, road traffic risk among others, causing many stakeholders to view these species as pests.

In this study we aimed to apply Bayesian areal disaggregation regression to ecological count data in the form of deer hunting bag returns collected by a government agency to produce trends of distribution and relative abundance, utilising this method for the first time in an ecological context. Using habitat classification data from publicly available remote sensing data, we aimed to map the changing distribution and relative abundance of deer in Ireland from 2000-2018. To evaluate the robustness of the predictions and demonstrate the reliability of Bayesian areal disaggregation regression to analyse regional count data in ecology, we challenged our model predictions by assessing their correlation with two independent datasets of deer distribution and relative abundance in Ireland using (a) predictions of deer distribution and relative abundance modelled by Morera-Pujol et al. (2022) using integrated species distribution models (therefore combining both presence-only and presence-absence data), and (b) faecal pellet group count data collected in field sites across Ireland.

## Methods

We deployed Bayesian areal disaggregation regression (Arambepola et al. 2021) to model the distribution and relative abundance of three species of deer (red, sika and fallow, respectively) in Ireland for the period 2000-2018. We used publicly available remote sensing data from the Corine Land Cover datasets (© European Union, Copernicus Land Monitoring Service, European Environment Agency) and publicly available hunting bag (also referred to as culling) returns collected yearly for each county of Ireland for each deer species (n = 26 counties, mean size: 2632.8km^2^, SD:_1618.4km^2^). We built 12 Bayesian areal disaggregation regression models to model the distribution of the 3 species for the years 2000, 2006, 2012, 2018, as CORINE data have been collected for these years, using the same methodology, allowing us to model the distribution and relative abundance of deer in Ireland over time. Culling returns were indeed also available since the 1990s, but we decided not to include them in the analysis due to the lack of consistent CORINE data for those years. All spatial and statistical analyses were conducted in R 4.2.0 (R Core Team, 2022).

### Environmental data and county-level deer culling returns

We used the CORINE habitat classification with some modifications: we excluded marine environments; we grouped all urban, industrial, and otherwise built up areas into the category “Artificial surfaces”; all bare rocks, sand beaches, burnt areas and other unvegetated habitats were merged into “Open spaces with little or no vegetation”; and we merged all agricultural land that was not pastures into “Planted vegetation and crops”. The 12 final habitat categories remaining were: planted vegetation and crops, pastures, natural grasslands, broad-leaved forest, coniferous forest, mixed forest, transitional woodland-shrub, moors and heathland, peat bogs, open spaces with little or no vegetation, inland waters and artificial surfaces. Using the vector layers for each resulting habitat, we calculated, for every pixel in a 5×5km raster, the proportion of the pixel covered by the habitat polygon. The resulting rasters then had pixel values between 0 and 1 representing the coverage of each habitat.

Yearly culling return data from hunters were obtained at a county level from the National Parks and Wildlife Service (NPWS). Data were submitted yearly by hunters using a form sent to NPWS by mail. The data consist of the number of harvested animals per county for red, sika, and fallow deer. Data were provided separately for males and females and, since 2015, juveniles, but all animals of each species were considered together for this analysis. In addition to the culling return data, we also obtained from the NPWS the number of licences issued yearly for each count. As the model was designed for use in epidemiology, it includes the option to account for human population given the logic that areas with higher population density are likely to have higher incidence rates of disease. Likewise, we included hunters (i.e., issued licences) as a proxy for population with the rationale that more hunters will likely lead to increased numbers of deer harvested. We thus capture the heterogeneous distribution of hunters throughout Ireland in our model to account for variability in culling returns. The raw data are depicted by species and year in Supplementary Material 1.

### Bayesian areal disaggregation regression models

We used the *disaggregation* package (Nandi et al. 2020) to run 12 Bayesian areal disaggregation models, one for each CORINE cycle (2000, 2006, 2012, 2018) and each prevalent deer species (red, sika and fallow).

We modelled the culling returns for each county (i) as a Poisson likelihood

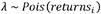

For each of the 26 counties (i), 5 x 5 km pixel (j) predictions are estimated using a weighted sum through the aggregation raster (a) that contains information on the number of hunters at a pixel level

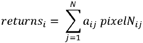

Within each county (i), the number of deer harvested per pixel (j) was modelled as

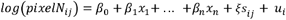

where β_0_ is the intercept, β _1_ to β _n_ are the effects of covariates X_1_ to X_n_, is a Gaussian Random Field (GRF) defined as ξ (*s*) ~ *N*(μ, ∑) with mean mμ and covariance matrix ∑.

We fit the models within the integrated nested Laplace approximation (INLA) framework (Lindgren and Rue 2015) which allowed us to model in continuous space using stochastic partial differential equations (SPDE, Lindgren et al. 2011). To do so we approximated continuous space through a triangular mesh over the study domain. We designed the spatial mesh to be 5×5km^2^, the specifications for the mesh were designed to ensure the predictive power of the model was robust, based on a sensitivity analysis by Arambepola et al. (2021) which concluded that polygons should consist of 200 pixels or less. Thus, we selected 5×5km^2^ to ensure that the majority of counties fit this criterion, only Cork (300 pixels), Mayo (246 pixels) and Galway (223 pixels) were above the recommended threshold of 200 pixels.

Priors were set on the parameters of the within county model. For the GRF, priors are set on the range, which controls the smoothness of the spatial field (i.e., the distance between peaks and troughs); and the variance, which controls the magnitude of these peaks and troughs. We used Penalised Complexity (PC) priors, a relatively new method that provides easily interpretable and controllable priors (Simpson et al. 2014) that are weakly informative, allowing the posterior distributions to be controlled mainly by the data. They penalise model complexity by forcing the model to its simplest realisation (the “base” model), a completely flat spatial field with infinite range and zero variance. The priors are specified by informing the model of “how far it is allowed to deviate” from the base model using the following specifications:

The prior on the range (ρ) is set providing the lower tail quantile and the probability so that

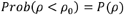

In this way we are limiting the range to values between infinite (the base model) and ρ_0_ with a probability of 95%. The prior on the variance is set on the standard deviation (sd), providing the upper tail quantile and the probability so that

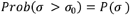

Thus, the sd value is limited between 0 (the base model) and sigma0 with a 95% probability.

### Model evaluation

For each of the 12 models (3 species x 4 years) we used the *predict()* function from the *disaggregation* package to obtain predicted deer density at a 5×5km resolution across Ireland. Since models were fitted in a Bayesian context through the *INLA* package, the prediction obtained at each location is not a point value but a distribution, from which we can produce the mean and corresponding 95% confidence intervals, thus obtaining a spatial estimate of the uncertainty of the prediction. Finally, to evaluate the performance of the model, we analysed the correlation (see below for full details) between the spatial predictions of our Bayesian areal disaggregation regression model and those from a concurrent analysis of deer distribution and relative density in Ireland, ran by Morera-Pujol et al. (2022); as a second and independent evaluation method, we analysed the correlation between the spatial predictions from our year 2018 models against independent field survey data capturing relative deer density collected in forest properties across Ireland in 2019. The methodologies used to obtain the data we challenged our predictions against are described below.

First, we used the predicted distributions for red, sika and fallow deer from Morera-Pujol et al. (2022), a first-time analysis of deer distribution and relative abundance in Ireland - though lacking the temporal dynamics we include here. These predictions were obtained by jointly modelling presence/absence and presence only data in an Integrated Species Distribution Model (ISDM). Data for the ISDM were obtained from different governmental agencies and citizen science initiatives and modelled using SPDE to include a random spatial effect in a Bayesian context, using the package *PointedSDMs*. To assess the performance of our disaggregation models against these, we calculated the Pearson correlation between the mean prediction of their models and the mean prediction of ours at the pixel level. We ran two computations, first we tested the correlation of relative abundance with our estimates of density at the pixel level, to evaluate how well our model predicted the abundance of deer per pixel. To do this we extracted the relative abundance from both model predictions and correlated them using the *cor.test* function in R. Secondly we tested the correlation of the spatial distribution of pixels where deer were predicted to be present, by overlaying the two raster files and correlating them using the *layerstats* function in the *raster* package to obtain pearsons correlation coefficient to evaluate how well our model predicted the distribution of the species per pixel compared to Morera-Pujol et al. 2022.

Secondly, we obtained faecal pellet group count density estimates from 2019 by an expert surveyor, full details of data collection are detailed in Morera-Pujol et al. (2022). All three species were target species for survey within 61 forest properties across Ireland. To challenge our model predictions, we extracted the mean prediction value for the pixel overlapping the forest surveyed, using the *extract()* function from the *raster()* package. We then calculated the Pearson correlation between this mean density and the estimated density from faecal pellet group count surveys for each species at each site.

## Results

The 12 Bayesian areal disaggregation models were fitted to the data with default priors for covariate effects, which resulted in small effect sizes with large uncertainty, and with the spatial field capturing most of the variability. We thus iteratively ran the models with increasingly informative priors, careful to balance the effect of the covariates and the spatial field for each species to ensure that the models were statistically robust while also making sense ecologically. For each species, we iteratively refined the model to find the most informative priors that were consistent across the four time periods so that the models could be comparable. We found best performance from, for the spatial field, a range of 50km to 85km and a sd of 0.5-2 across the three species and values between 1.51 and 3 for the slope of the fixed effects The priors and their corresponding posterior mean and standard deviation are shown in Table 1 for the 2018 model as an example, full prior and posterior range for all the models are shown in Supplementary Material 2 and the slopes of the fixed effects are shown in Supplementary Material 3..

The spatial distribution and relative density of Ireland’s three prevalent deer species (red, sika and fallow) as predicted by our models for the years 2000, 2006, 2012 and 2018 show that much of the territorial expansion of all three deer species originate from high density population centres with red deer prevalent in the north west of the country, sika deer in the south west and the east, and fallow deer in the midlands of Ireland. Over the study period we observe a low-density proliferation of all three populations into novel territory with increasing density in the original populations (Fig. 1) with the upper and lower 95% confidence intervals for the distributions of each species for each year in Supplementary Material 4. These trends are shown dynamically in species specific maps showing population growth and territorial expansion in .gif format presented in Supplementary Material 5 (red deer), 6 (sika deer) and 7 (fallow deer).

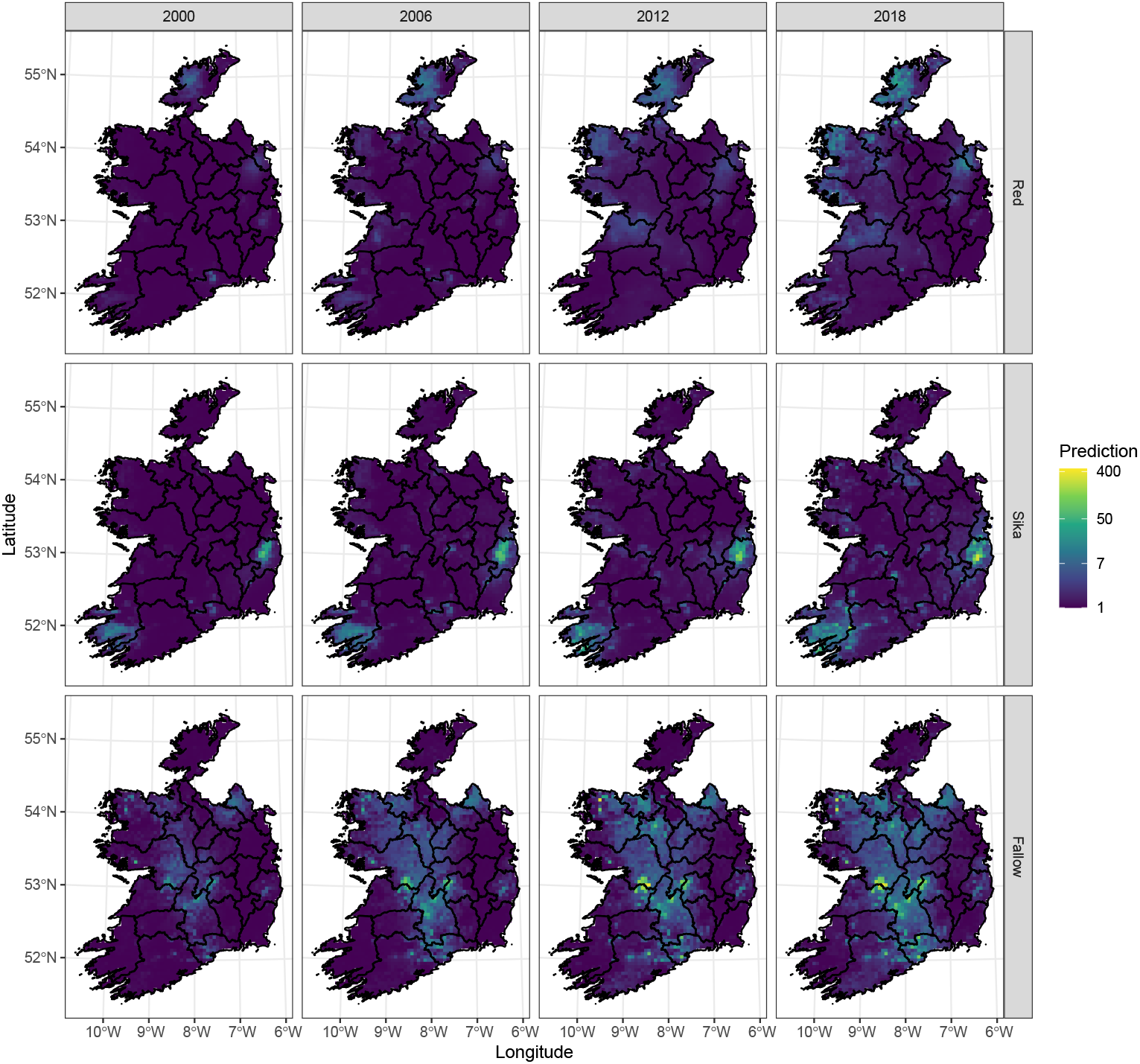

We observed changes in habitat preferences over the study period for each of the three species, with noticeable shifts from natural to modified landscapes. For red deer, we found a switch in selection over the study period from natural grassland habitat to pasture and peat bogs, however, there is also a relative decrease in the selection of planted vegetation (Figure 2, top row). For sika deer we detected an increased reliance on conifer forestry, heathland and planted vegetation, while the effect of peat bog was reduced over the study period (Figure 2, middle row). For fallow deer we found that there was a relative decrease in the use of pasture and peat bog habitats and increasingly fallow deer selected for artificial surfaces and planted vegetation during the period 2000-2018 (Figure 2, bottom row).

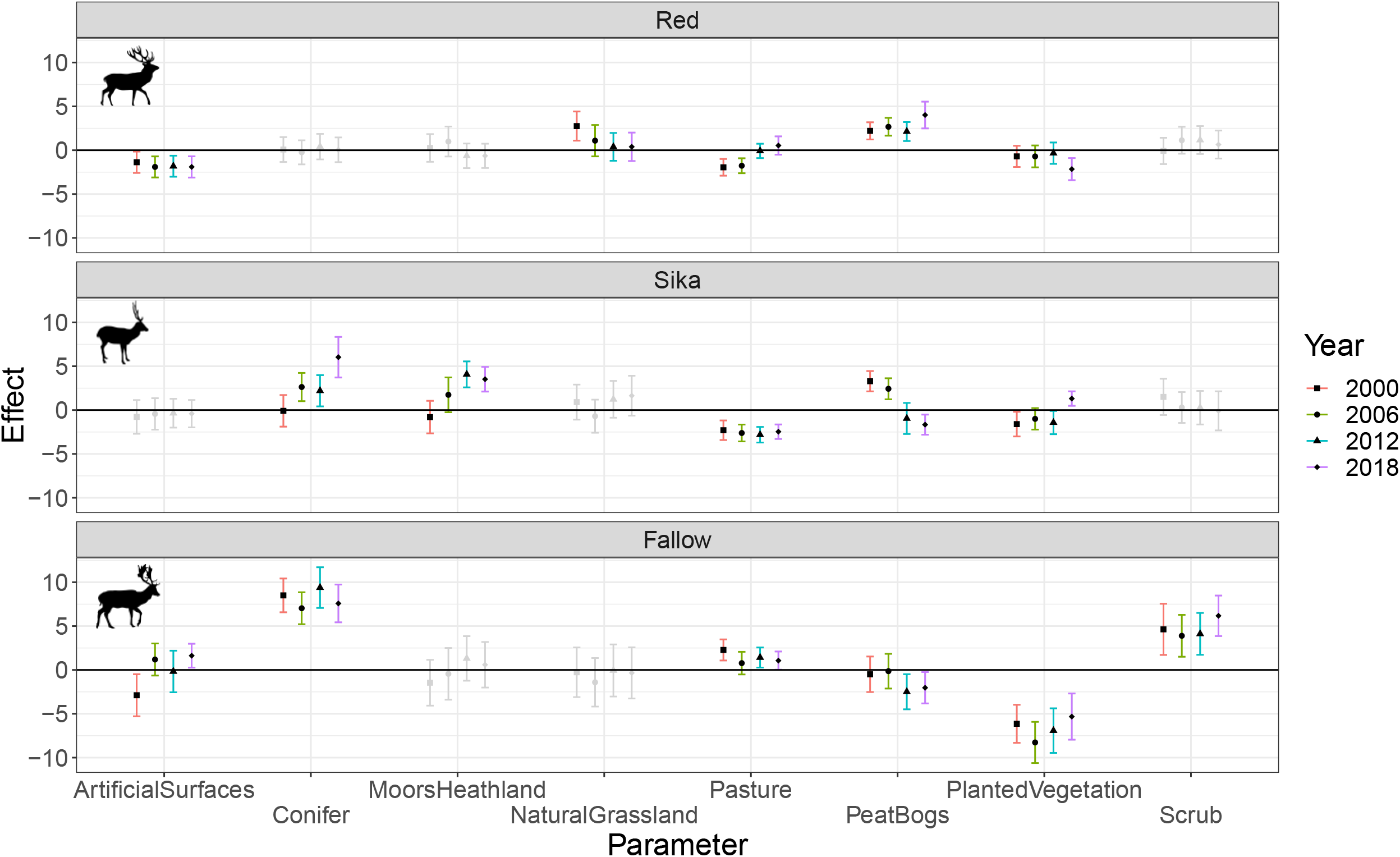

We found that our model predictions were highly correlated with the estimates provided by Morera-Pujol et al. (2022) and with field survey density estimates from 61 sites across Ireland. We compared Morera-Pujol et al. (2022) estimates of deer abundance at the pixel level with our model predictions for relative density and found correlation coefficients of 0.76 for red deer, 0.60 for sika deer and 0.79 for fallow deer. For spatial distribution overlap between our model predictions and Morera-Pujol et al. (2022) we obtained correlation coefficients of 0.69 for red, 0.70 for sika and 0.59 for fallow, respectively. Finally, when comparing the relative density we predicted at the pixel level against real world field data collected by an expert surveyor, we found that our model performed strongly, obtaining correlation coefficients of 0.69 for red deer, 0.68 for sika deer and 0.40 fallow deer against the pellet group count field data.

We compared the spatial distributions as predicted by an INLA approach and our Bayesian areal disaggregation regression approach in Figure 3 to demonstrate the spatial predictions of the two methodologies despite vast differences in data quantity and quality, with over 20 years of point data collection for the INLA approach and one year of poor resolution areal data for each Bayesian areal disaggregation regression model. We visualised the real-world distribution and abundance of hunting bag data we collected at the county level for 2018 and compared that with the prediction of spatial distribution and relative density per 5×5km^2^ grid cell we predict in the model for each of the three species in Figure 4.

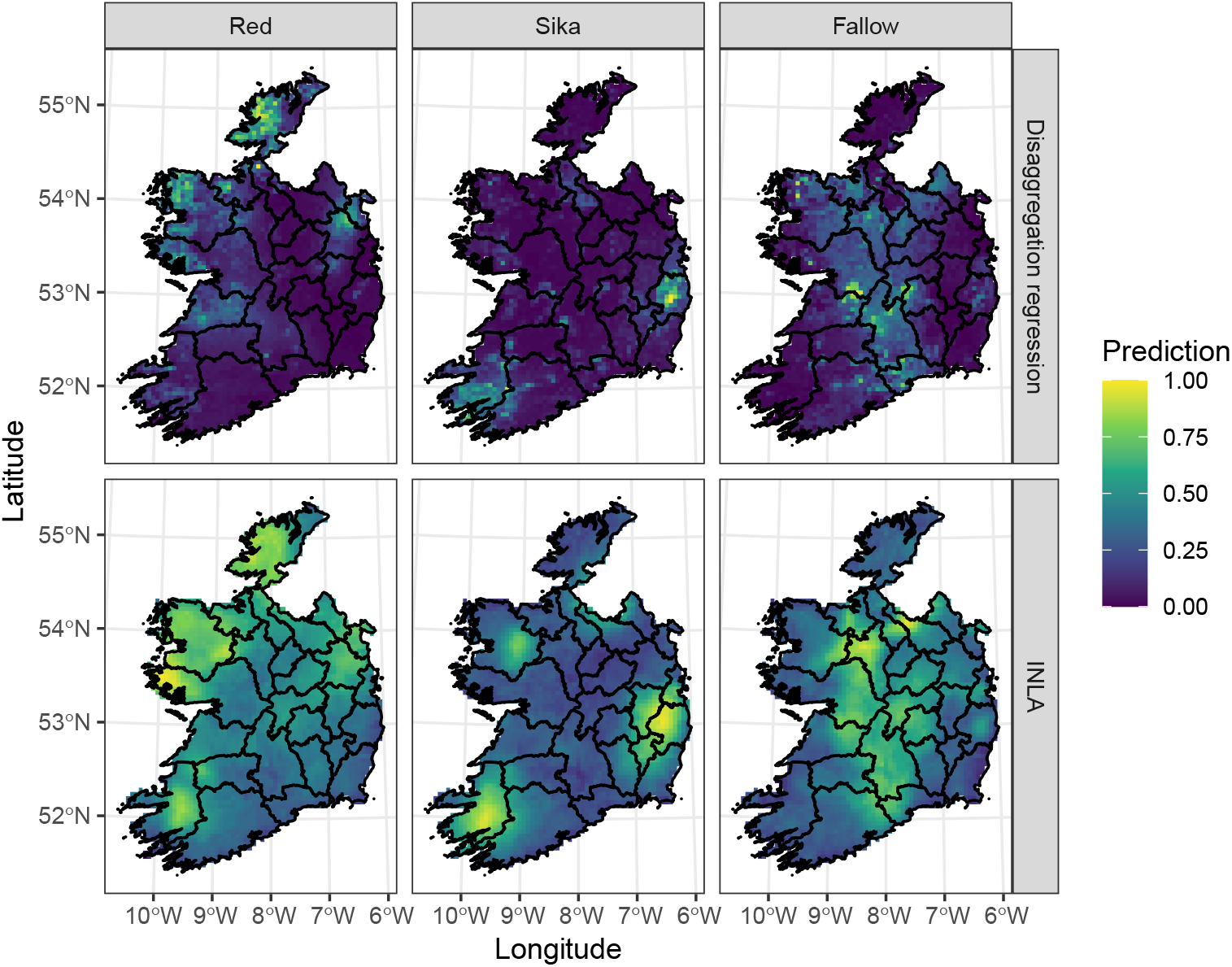

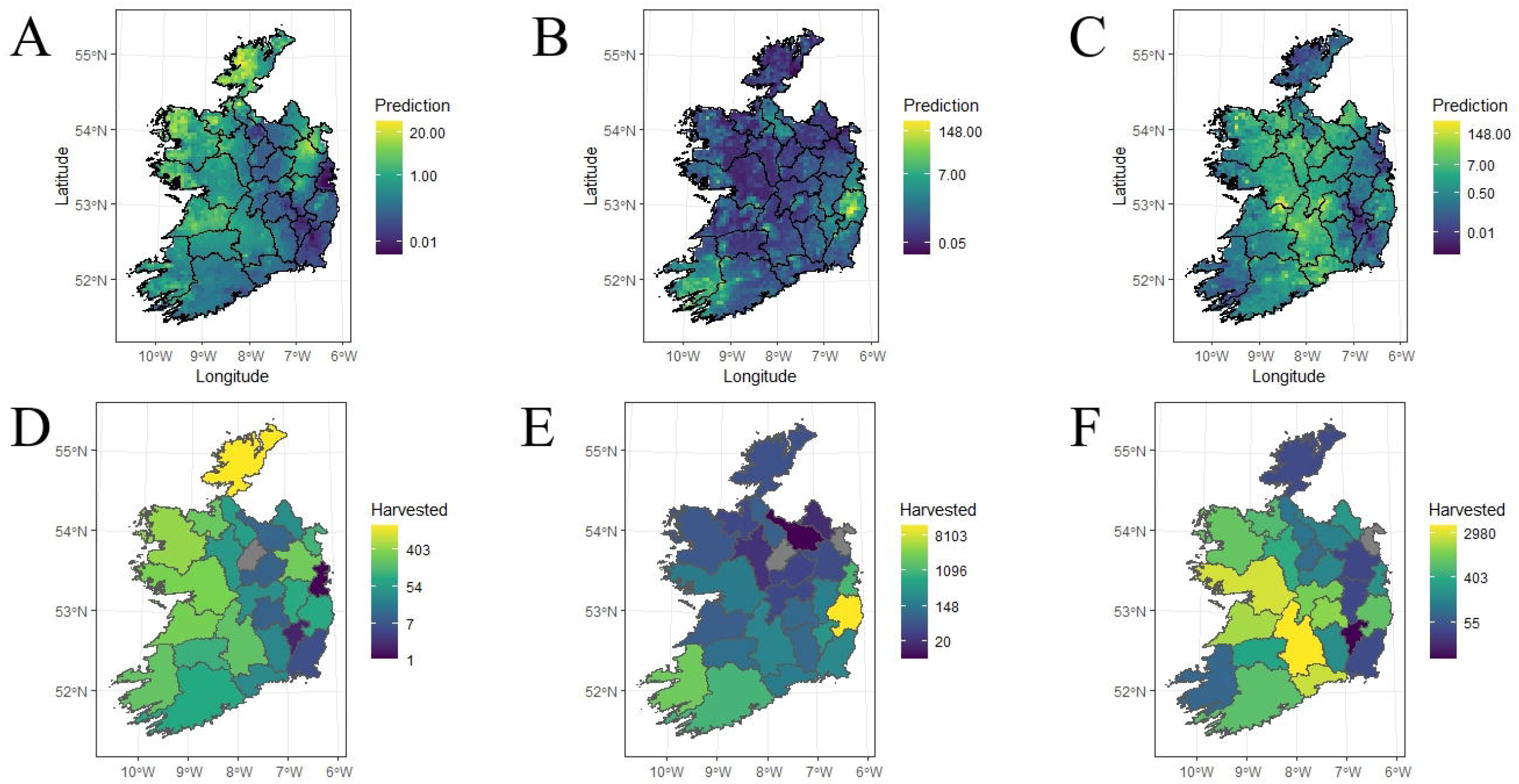

## Discussion

### Bayesian areal disaggregation regression: a new tool for wildlife conservation and management

In this study we demonstrated that Bayesian areal disaggregation regression can be used to predict high resolution wildlife distribution and relative abundance from low resolution count data. We evaluated the predictions of our model with two independent estimates of deer distribution and relative abundance in Ireland, finding high correlation with our model estimates. The strength of our method compared to traditional techniques is that we produced high-resolution, statistically robust and trustworthy estimates of multi-species distribution and relative density using low-quality count data collected in a labour-free and inexpensive manner. In doing so, we have introduced a new method in applied ecology to produce distribution and relative abundance data for wildlife and provided a new analytical framework for low-resolution areal data which may previously have been unexploited. These findings allow for change in the wildlife management pipeline (from data collection to informed decision making), opening new scenarios to exploit previously unused dark data from wildlife counts to produce high-resolution indices of distribution and abundance, saving resources for more periodic high-resolution wildlife monitoring. Our analysis agrees with and provides a new methodology that bolsters the findings of Boyce et al. (2012) that argues for cost-effective methods to analyse count data from hunting bag or wildlife observations at areal level. Bayesian areal disaggregation regression is a robust choice that can be used to analyse such data, providing population metrics comparable to expensive high-resolution surveys (e.g., aerial surveys). Periodic high-resolution surveys could be undertaken to evaluate model predictions, thus saving resources and allowing for the cost-effective and science-based management of wildlife populations “by the seat of your pants” (*sensu* Boyce et al 2012).

Understanding the effects of land-use change and human perturbation on wildlife population ecology is essential to design data-driven decision-making programmes to manage wildlife as land-use continues to change in line with social, economic and political landscapes (Renwick et al. 2013; O’Rourke, 2019; Lecuyer et al. 2022). Conflict species such as deer that have successfully navigated human modification and expanded out of their historic range, must be studied to understand and prepare for future land-use change scenarios e.g., increased afforestation, farm abandonment, rewilding and continued encroachment of human settlements that have been observed worldwide (Hoogendorm et al. 2019; Castillo et al. 2021). Information on relative population size and distribution is essential to the conservation and management of wildlife and can form the foundation of higher-resolution research programmes to further refine management objectives and decision-making protocols (Gillson et al. 2019; Morera-Pujol et al. 2022). Where high-resolution data are not available to analyse, it may be necessary to use lower-resolution data to obtain baseline information on species distribution and relative population density.

The method we use in this study allows for low-resolution areal count data, in this case Ireland’s county-level national hunting bag returns, to produce high-resolution information on wildlife population dynamics. We use this approach to fill an important research gap across a national landscape, however, this methodology can inform more granular scales depending on the spatial resolution of areal count data. Hunting bag data is commonly collected in countries where hunting is nationally coordinated by the state, allowing this method to be transposed across species (e.g., avian game and waterfowl, mammalian predators and ungulates), and countries groups where hunting bag data are collected (Apollonio et al. 2010). Additionally, alternative forms of count data (e.g., citizen science counts) collected at areal level can be utilised to produce baseline information for management at the scale the data is collected at. Furthermore, advancements in remote sensing equipment means that high-resolution environmental data is obtainable globally through publicly available programmes e.g., Google Earth Engine. The relatively low data demands for this methodology to produce high resolution predictions for species range and relative abundance should ensure that this method becomes established in applied ecology to provide baseline information (e.g., species distribution) for species that are counted at areal level and but lack fine-scale systematic monitoring schemes.

Bayesian areal disaggregation regression has proved to be a powerful tool for wildlife biologists to exploit low-resolution data to produce high-resolution predictions of distribution and relative density, however, there are challenges associated with the method that must be disclosed. The creation of the spatial mesh requires careful specification in all models requiring one. In the case of the disaggregation regression models and following Arambepola et al. (2021) we created a mesh that resulted in an appropriate number of triangles per aggregation unit. of the size of the triangles and complexity of the boundaries, among others. Another sensitive part of setting up these models is the prior specification. This requires an iterative approach, testing different prior values and evaluating the posterior distributions and model predictions. By comparing the priors set to the range and SD of the spatial field to their posterior distributions, we can iteratively modify the priors to make sure they are not pulling the model too far away from the data space. The size of the covariate effects can be controlled using the priors for the slope (mean and SD of a Gaussian distribution). By adjusting the priors of both the spatial field and the covariate effects we can balance the effects of the spatial field and the covariate effects to produce statistically robust and ecological significant model predictions. The obtention of full posterior distributions for the spatial prediction allows us to obtain measures of uncertainty (e.g., SD or 95% CI), which can help diagnose inaccuracies in the model. Perhaps most important for the use of these models in wildlife conservation and management is the validation of model predictions with real world estimates of distribution and density. While our models produced outstanding correlations with real world data, the absence of these validation datasets would severely impact our ability to trust the output of the model. This method can be used by institutions seeking to exploit dark-data that has previously gone unused due to concerns about the spatial resolution of the data, however it is imperative that these models are not used in isolation, but that they are routinely correlated against high-resolution survey data to ensure they are robust and capable of being used in a conservation and management decision making framework, (*sensu* Boyce et al. 2012).

### Lesson learnt from the Irish study case

Of the three study species included in this study only red deer are native to Ireland, and within that population, research suggests that only the Killarney, Co. Kerry (Fig. 1 - southwest hotspot) subpopulation in the south-west of the country is authentically native to Ireland (McDevitt et al. 2009). It is likely that human introductions are the strongest driver of deer distribution in Ireland, with much of the range expansion we observe radiating out of founder populations in proximity to their historical introductions i.e. Kerry, Galway (Fig. 1 - west), Meath (Fig. 1 - east), and Donegal (Fig. 1 - northwest), for red deer (McDevitt et al. 2009; Carden et al. 2011), Kerry and Wicklow (for sika deer (McDevitt et al. 2009), and historical sites of Anglo-Norman castles and deer parks located in Ireland’s east and midland counties for fallow deer (Beglane et al. 2018). It is also possible that there have been many local introductions in the 20th and 21st century that have not been documented in the literature but may have contributed to range expansion or population growth for deer in Ireland.

Ireland is representative of many countries across the world where deer population growth and range expansion has occurred since the mid-20th century, typified by predator eradication, increased resources due to agriculture and afforestation (Coté et al. 2004). Ireland is devoid of large predators capable of hunting deer since the extinction of brown bear (*Ursus arctos*) and lynx (*Lynx lynx*) following the last glacial maximum, and the extermination of the wolf (*Canis lupus*) in the 18th century, which had previously been widespread throughout Ireland (Sleeman, 2008; Hickey, 2000). Predator persecution has globally led to irruptions in deer populations and range expansion associated with increased human-wildlife conflict and ecological degradation (Ripple et al. 2010; Ripple et al. 2012). Ireland has followed a state policy of afforestation since 1950, increasing total forestry area from 2.1% to 11%, covering 770,020 hectares of total land-area in 2020 (DAFM, 2020). Concurrently, increasing forest cover and agricultural cover provide novel cover and resources which appear to drive the population growth and range expansion we observed in our study. Thus, Ireland faces a suite of management challenges that arise when deer are overabundant e.g. forest and crop damage (Pérez-Espona et al. 2009; Murphy et al. 2013), agricultural disease such as bovine tuberculosis (Crispell et al. 2020; Kelly et al. 2021), human health and safety (Gray et al. 1996; Langbein et al. 2010) and animal welfare (Tranulis et al. 2021). Challenges that cannot be managed effectively without an evidence-based approach.

We detected an increased preference for modified landscapes in lieu of natural ones across all three species. Red deer switched from natural grasslands to grazing pastures over the study period, potentially placing them in contact with agricultural animals and providing pathways for disease transmission, such as bovine tuberculosis. Conifer forest and moors became increasingly attractive to sika deer, which could result in damage to commercial forestry. This coincided with a decreasing selection/preference of peat bogs, though this may be confounded by the afforestation of peat bogs with conifer forestry (Black et al. 2008). Fallow deer increasingly selected artificial surfaces (i.e., human settlement and infrastructure), and have consistent preference for conifer forest over the study period which may also place this species close to the human-wildlife conflict interface. While land-use change seems to predominantly drive deer expansion and population growth, there are existing confounding factors which we hypothesise may also influence these trends, for instance, the hunting model in Ireland.

We hypothesise that within high density “hot spots” hunting pressure is high, resulting in displacement of deer into surrounding areas where conflicts arise, which then triggers the approval of “out of season” licences for pest species that causes a year-round low-intensity hunting pressure that pushes deer into surrounding available habitats. Research conducted in Scandinavia detected behavioural responses of red deer to human hunting pressure whereby red deer adjusted their spatial ecology increasing the likelihood of migration on the first day of the hunting season (Meisinget et al. 2022). If occurring in Ireland, this effect may potentially be creating complex source-sink dynamics that are contributing to the range expansion we observe, though confirming this will require more granular research using wildlife tracking devices. Furthermore, human-mediated ecological disturbances (e.g., clearfelled forestry, construction, agricultural activities etc) may also play a role in manipulating species spatial ecology, behaviour, and population dynamics that should also be examined further (Mysterud et al. 2020; Murphy et al. 2022). Thus, to achieve sustainable deer management and healthy populations it is important to understand the distribution and relative density of the species at landscape scale, to highlight high-density source populations and potential sink habitats in the vicinity to allocate management efforts accordingly (Wäber et al. 2013).

### Conclusions

In this study we used low-resolution areal count data to model deer range expansion and population growth, subsequently evaluating these findings with real-world monitoring data, finding high correlations between model prediction and survey data. These results clearly demonstrate the applicability of Bayesian areal disaggregation regression for applied ecological research. Furthermore, these models provide inference on the relative effect size with 95% confidence intervals of each covariate which is essential for designing informed science-based management of wildlife, their habitats and their interface with human populations. Our analysis provides distribution and population trends for three species of deer across a national landscape to fill an important knowledge gap in Ireland, however, this methodology can be applied to any species and spatial scale where areal count data are collected provided that there is covariate information available and standardised jurisdictional division between the counts. Certainly, this method can now be confidently considered within the applied ecological toolbox for analysing areal count data.

Wildlife monitoring to understand species distribution and population dynamics will continue to be a priority for research and management in applied ecology, especially in the case of conflict species and species of conservation concern (Moussy et al. 2021). The collection of high-resolution wildlife data, together with environmental remote sensing data, capable of being analysed to inform management decisions is a costly and effort intensive task (Marvin et al. 2016). Poor practice in data sharing and curation can reduce the impact of previous monitoring efforts if dark data remains stored and unused in repositories collected by previous users (Hampton et al. 2013). Applied ecologists should be adaptable and prepared to rescue dark data with new statistical methods which allow low resolution data to be repurposed for use in novel analytical frameworks to contribute to research and management (Karlsson, 2022). As data science methodologies begin to develop across disciplines and novelty emerges from improved computational power, ecologists must continue to test their efficacy within our own research programmes and evaluate their performance on the data we collect. In an era of extensive anthropogenic land-use change, human-wildlife conflict, worsening biodiversity loss and global climate change, scientists working on these challenges can harness increasingly powerful tools and data to understand and manage wildlife populations and their habitats in a changing world.

## Data Availability Statement

All data will be uploaded to Dryad repository

## Contributions Statement

Conceptualization: K.J.M, S.C., V.M.-P; Methodology: K.J.M.,, V.M.P; Formal analysis: K.J.M.; Writing—original Draft: K.J.M.; Writing—review and editing: K.J.M., S.C., V.M.-P., T.B; Visualisation: K.J.M., V.M.-P.; Supervision: V.M.-P., S.C.

