## Supplementary material for "Bayesian areal disaggregation regression to predict wildlife distribution and relative density with low-resolution data": Table 2

Table 2. Pearsons correlation coefficient for model predictions when correlated against independent data. Relative abundance refers to the correlation between our model prediction of relative density with relative abundance estimates from Morera-Pujol et al. 2022. Spatial overlap abundance refers to the correlation between the spatial overlap of species distribution at the pixel level between our model prediction and the distribution computed by Morera-Pujol et al. 2022. Finally, we correlated our model predictions of species density and distribution with field data in the form of faecal pellet group counts from 61 sites across Ireland.

| **Species** | **Relative abundance**  **Morera-Pujol et al. 2022 prediction** | **Spatial overlap**  **Morera-Pujol et al. 2022 prediction** | **Faecal pellet group count density estimates** |
| --- | --- | --- | --- |
| Red deer | 0.76 | 0.69 | 0.69 |
| Sika deer | 0.60 | 0.70 | 0.68 |
| Fallow deer | 0.79 | 0.59 | 0.40 |
