## Supplementary figures and images for "Bayesian areal disaggregation regression to predict wildlife distribution and relative density with low-resolution data"

### Supplementary Material 1

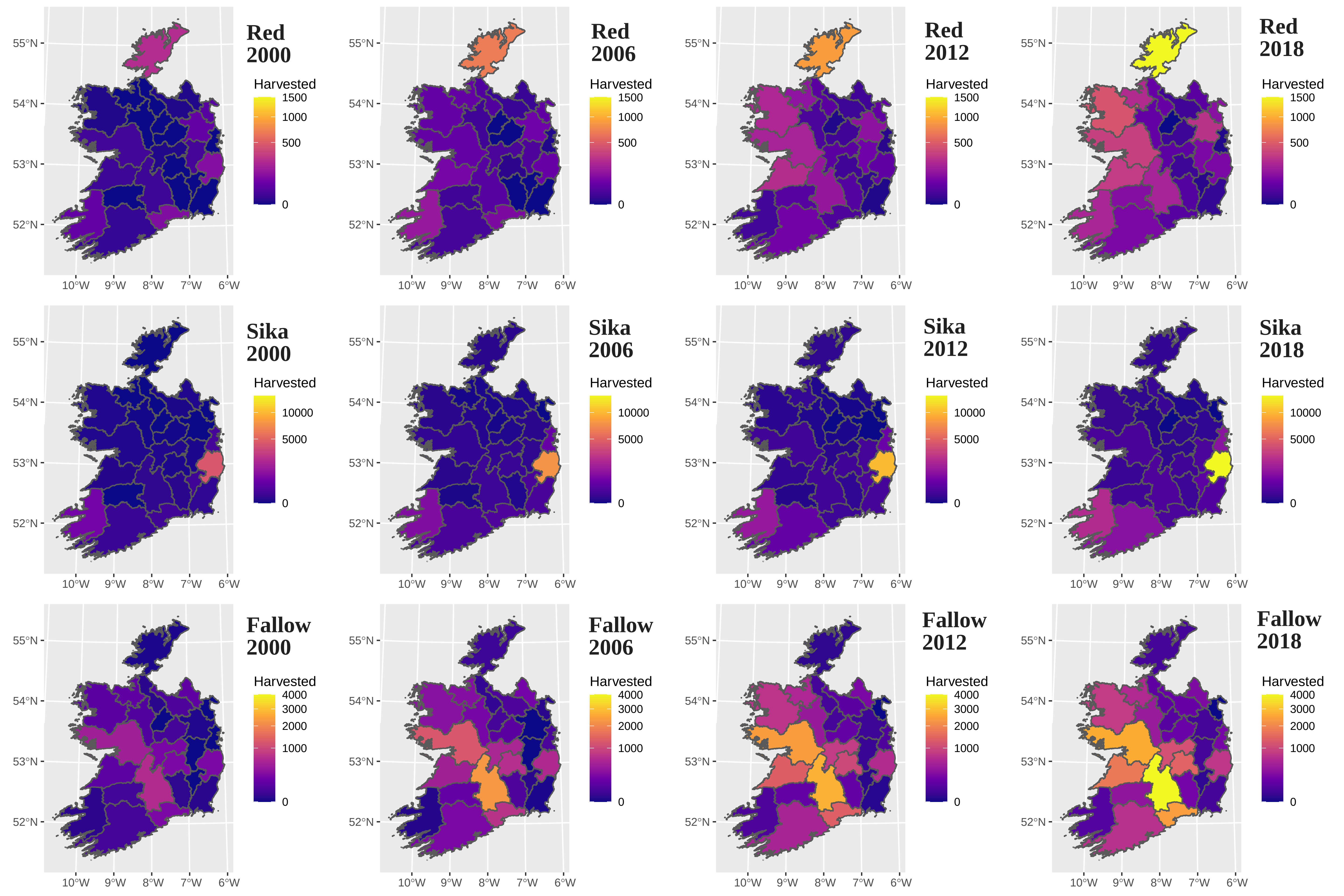

### Supplementary Material 4

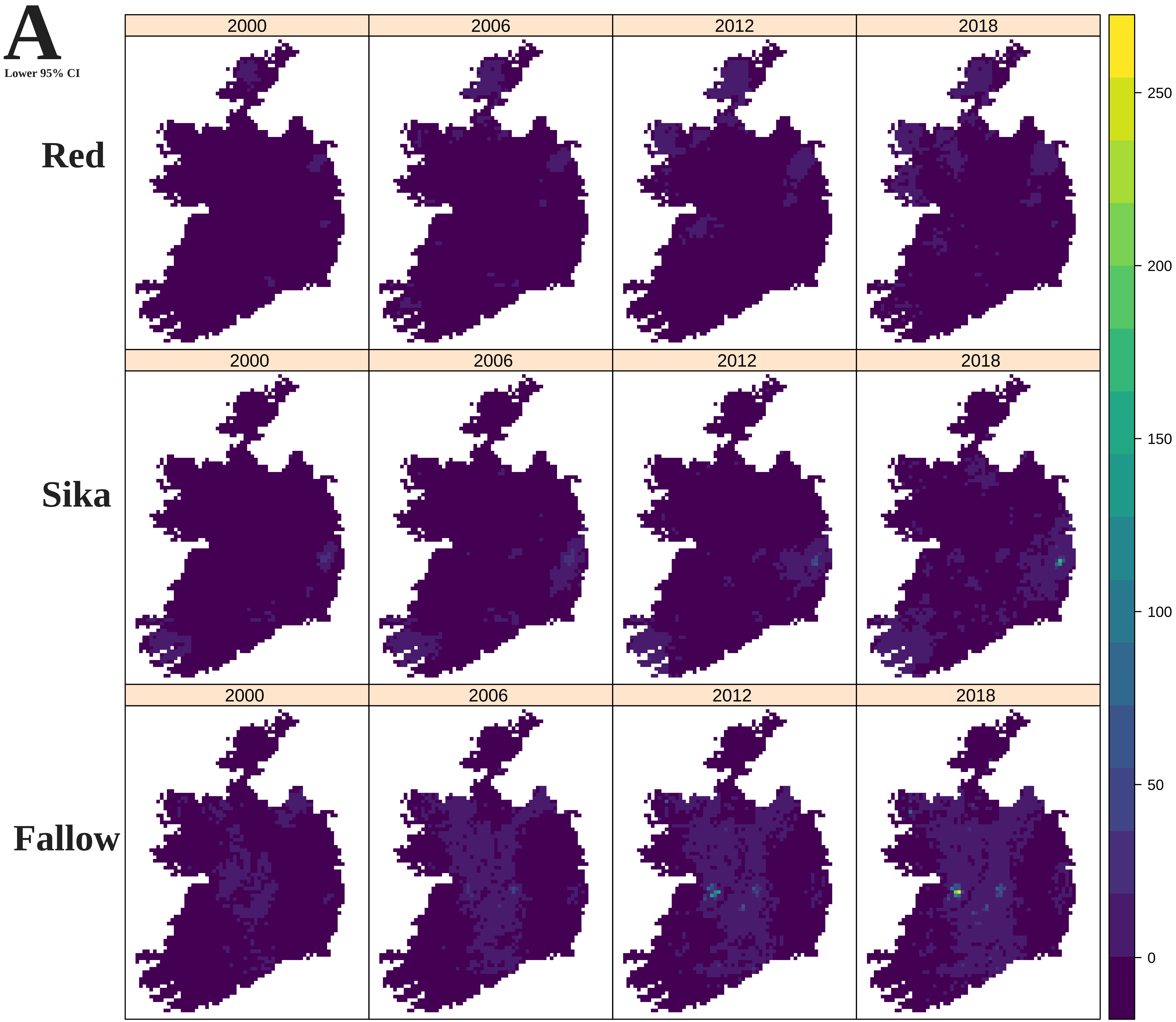

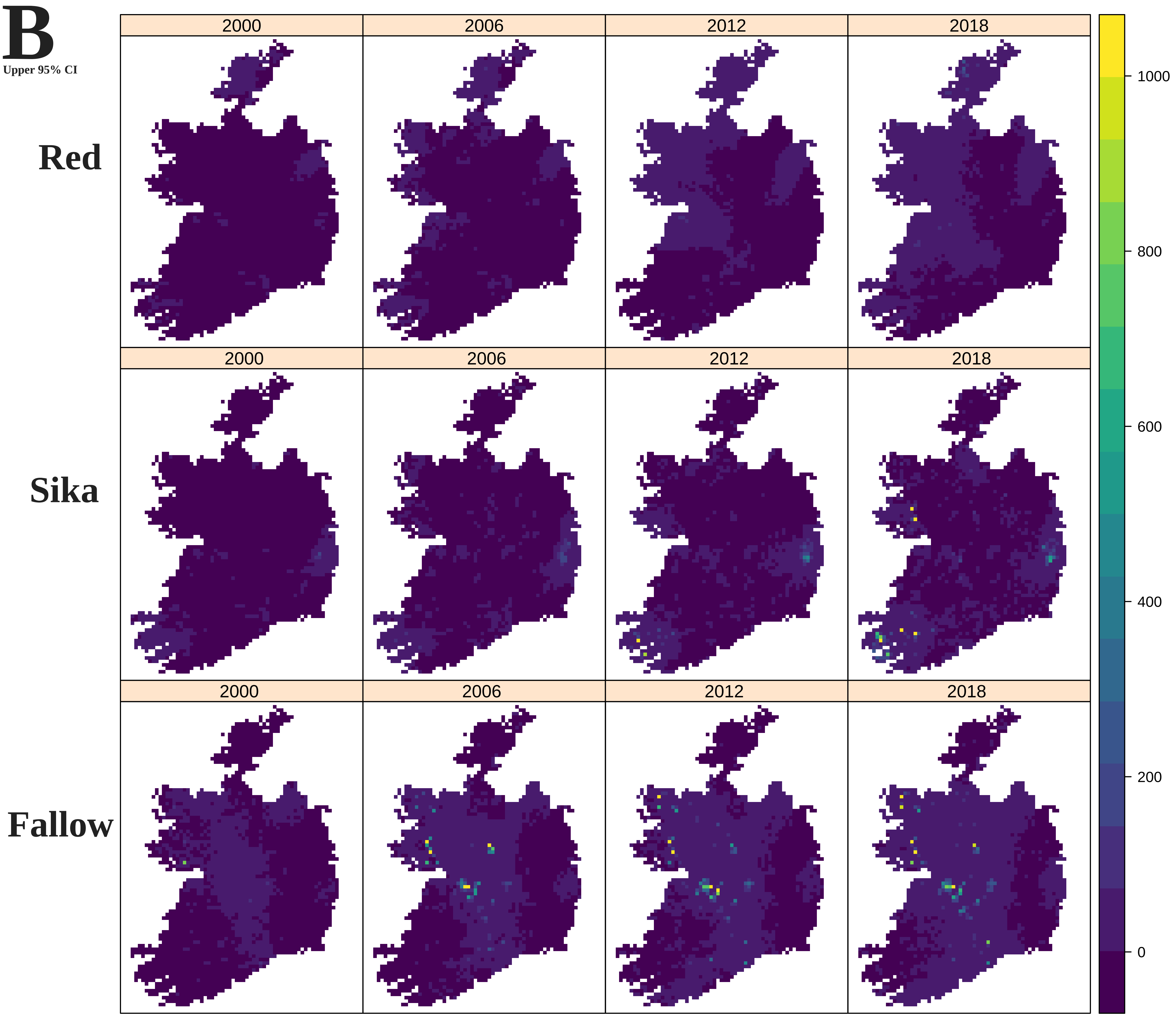

### Supplementary Material 5

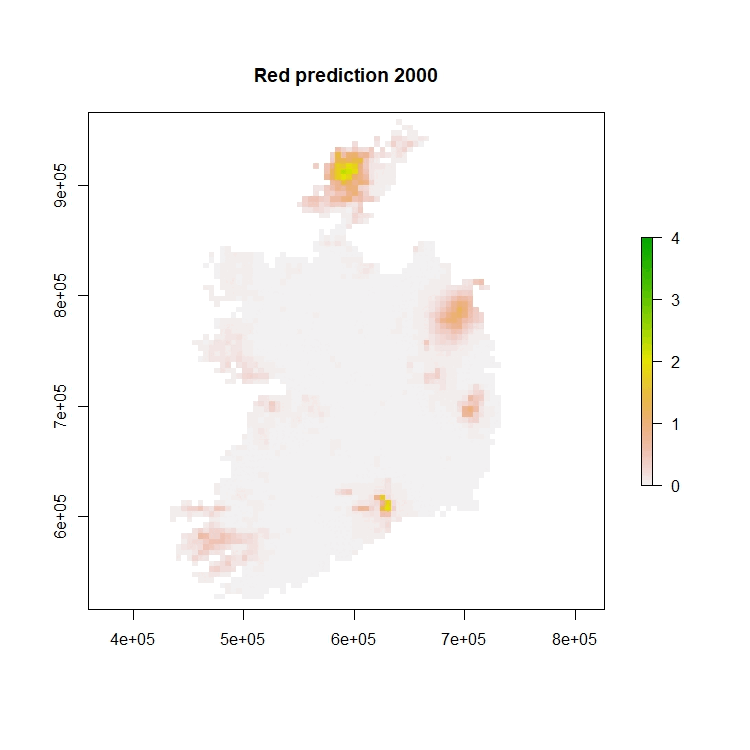

### Supplementary Material 6

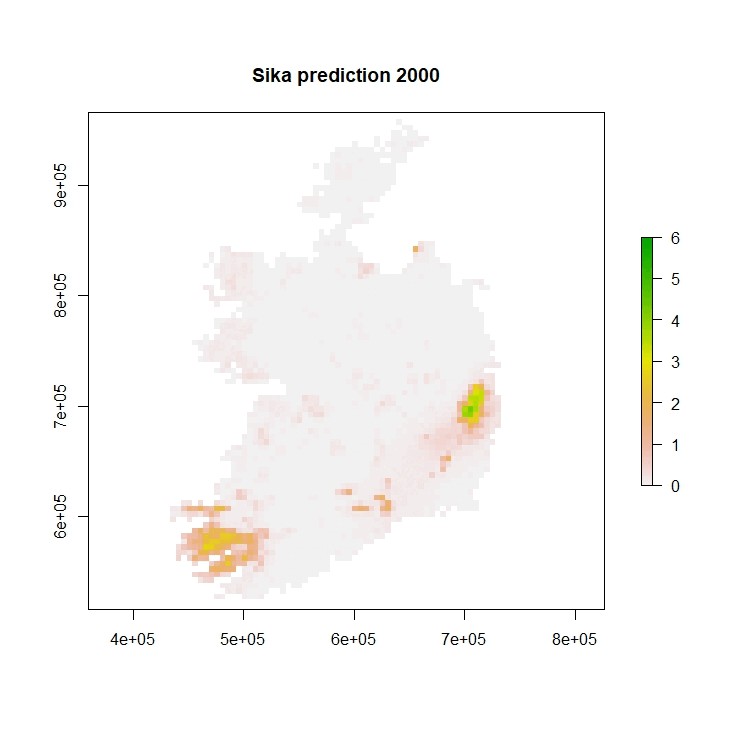

### Supplementary Material 7

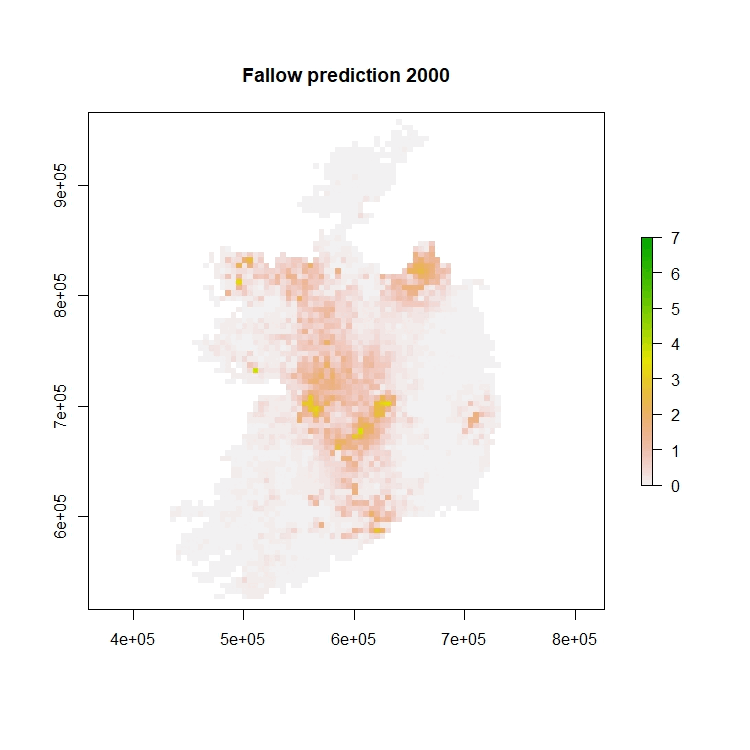
