## Supplementary Material 2 for "Bayesian areal disaggregation regression to predict wildlife distribution and relative density with low-resolution data"

Supplementary Material 2A: Priors' specification and posterior distribution estimated for all the parameters of the three species-specific (red, sika and fallow deer) models for 2000.. Spatial fields are defined in our model by their range (ρ) and standard deviation (σ). Priors are set on the range and standard deviation as penalised complexity (PC) priors and so the lower limit for the range and the upper limit for the SD are specified, together with a probability value for each. The prior to the intercept of the model was set following Arambepola et al. (2021). The fixed effects are defined by their slopes ($\beta_{i})$, and to these we set priors as a Gaussian distribution defined by its mean and SD. The posterior distributions of all parameters were obtained from the model and are described here by their mean and standard deviation. * The slopes of the fixed effects are shown in Supplementary Material 2b.

| **Species** | **Parameter** | **Prior** | **Posterior mean** | **Posterior SD** |
| --- | --- | --- | --- | --- |
| Red deer | Unstructured random effect | **-** | **-** | **-** |
|  | Range of the spatial field (m) | ρ_0_ = 50000, P(ρ) = 0.05 | 54720.85 | 0.15 |
|  | Standard deviation of the spatial field | σ_0_ = 0.5, P(σ) = 0.1 | 1.08 | 0.15 |
|  | Intercept | $\beta_{0} \sim N\left( -4 , 4 \right)$ | -3.28 | 0.67 |
|  | Slopes of the fixed effects | $\beta_{i} \sim N(0.0, 1.51)$ | SM2b | SM2b |
| Sika deer | Unstructured random effect | **-** |  |  |
|  | Range of the spatial field (m) | ρ_0_ = 85000, P(ρ) = 0.05 | 81633.91 | 0.15 |
|  | Standard deviation of the spatial field | σ_0_ = 0.1, P(σ) = 0.1 | 1.04 | 0.08 |
|  | Intercept | $\beta_{0} \sim N\left( -4 , 2 \right)$ | -2.99 | 0.83 |
|  | Slopes of the fixed effects | $\beta_{i} \sim N(0.0, 2)$ | SM2b | SM2b |
| Fallow deer | Unstructured random effect | **-** |  |  |
|  | Range of the spatial field (m) | ρ_0_ = 75000, P(ρ) = 0.05 | 63576.55 | 0.13 |
|  | Standard deviation of the spatial field | σ_0_ = 2, P(σ) = 0.1 | 1.06 | 0.08 |
|  | Intercept | $\beta_{0} \sim N\left( -4 , 3 \right)$ | -3.38 | 1.01 |
|  | Slopes for the fixed effects | $\beta_{i} \sim N(0.0, 3)$ | SM2b | SM2b |

Supplementary Material 2B: Priors' specification and posterior distribution estimated for all the parameters of the three species-specific (red, sika and fallow deer) models for 2006. Spatial fields are defined in our model by their range (ρ) and standard deviation (σ). Priors are set on the range and standard deviation as penalised complexity (PC) priors and so the lower limit for the range and the upper limit for the SD are specified, together with a probability value for each. The prior to the intercept of the model was set following Arambepola et al. (2021). The fixed effects are defined by their slopes ($\beta_{i})$, and to these we set priors as a Gaussian distribution defined by its mean and SD. The posterior distributions of all parameters were obtained from the model and are described here by their mean and standard deviation. * The slopes of the fixed effects are shown in Supplementary Material 2b.

| **Species** | **Parameter** | **Prior** | **Posterior mean** | **Posterior SD** |
| --- | --- | --- | --- | --- |
| Red deer | Unstructured random effect | **-** | **-** | **-** |
|  | Range of the spatial field (m) | ρ_0_ = 50000, P(ρ) = 0.05 | 54720.85 | 0.15 |
|  | Standard deviation of the spatial field | σ_0_ = 0.5, P(σ) = 0.1 | 0.94 | 0.09 |
|  | Intercept | $\beta_{0} \sim N\left( -4 , 4 \right)$ | -2 | 0.15 |
|  | Slopes of the fixed effects | $\beta_{i} \sim N(0.0, 1.51)$ | SM2b | SM2b |
| Sika deer | Unstructured random effect | **-** |  |  |
|  | Range of the spatial field (m) | ρ_0_ = 85000, P(ρ) = 0.05 | 78433.00 | 0.17 |
|  | Standard deviation of the spatial field | σ_0_ = 0.1, P(σ) = 0.1 | 0.93 | 0.08 |
|  | Intercept | $\beta_{0} \sim N\left( -4 , 2 \right)$ | -1.47 | 0.75 |
|  | Slopes of the fixed effects | $\beta_{i} \sim N(0.0, 2)$ | SM2b | SM2b |
| Fallow deer | Unstructured random effect | **-** |  |  |
|  | Range of the spatial field (m) | ρ_0_ = 75000, P(ρ) = 0.05 | 76879.92 | 0.13 |
|  | Standard deviation of the spatial field | σ_0_ = 2, P(σ) = 0.1 | 1.09 | 0.13 |
|  | Intercept | $\beta_{0} \sim N\left( -4 , 3 \right)$ | -1.8 | 1.09 |
|  | Slopes for the fixed effects | $\beta_{i} \sim N(0.0, 3)$ | SM2b | SM2b |

Supplementary Material 2C: Priors' specification and posterior distribution estimated for all the parameters of the three species-specific (red, sika and fallow deer) models for 2012.. Spatial fields are defined in our model by their range (ρ) and standard deviation (σ). Priors are set on the range and standard deviation as penalised complexity (PC) priors and so the lower limit for the range and the upper limit for the SD are specified, together with a probability value for each. The prior to the intercept of the model was set following Arambepola et al. (2021). The fixed effects are defined by their slopes ($\beta_{i})$, and to these we set priors as a Gaussian distribution defined by its mean and SD. The posterior distributions of all parameters were obtained from the model and are described here by their mean and standard deviation. * The slopes of the fixed effects are shown in Supplementary Material 2b.

| **Species** | **Parameter** | **Prior** | **Posterior mean** | **Posterior SD** |
| --- | --- | --- | --- | --- |
| Red deer | Unstructured random effect | **-** | **-** | **-** |
|  | Range of the spatial field (m) | ρ_0_ = 50000, P(ρ) = 0.05 | 59874.14 | 0.14 |
|  | Standard deviation of the spatial field | σ_0_ = 0.5, P(σ) = 0.1 | 1.15 | 0.08 |
|  | Intercept | $\beta_{0} \sim N\left( -4 , 4 \right)$ | -1.91 | 0.66 |
|  | Slopes of the fixed effects | $\beta_{i} \sim N(0.0, 1.51)$ | SM2b | SM2b |
| Sika deer | Unstructured random effect | **-** |  |  |
|  | Range of the spatial field (m) | ρ_0_ = 85000, P(ρ) = 0.05 | 79221.26 | 0.15 |
|  | Standard deviation of the spatial field | σ_0_ = 0.1, P(σ) = 0.1 | 0.81 | 0.09 |
|  | Intercept | $\beta_{0} \sim N\left( -4 , 2 \right)$ | -0.35 | 0.71 |
|  | Slopes of the fixed effects | $\beta_{i} \sim N(0.0, 2)$ | SM2b | SM2b |
| Fallow deer | Unstructured random effect | **-** |  |  |
|  | Range of the spatial field (m) | ρ_0_ = 75000, P(ρ) = 0.05 | 77652.58 | 0.08 |
|  | Standard deviation of the spatial field | σ_0_ = 2, P(σ) = 0.1 | 1.01 | 0.13 |
|  | Intercept | $\beta_{0} \sim N\left( -4 , 3 \right)$ | -1.66 | 1.02 |
|  | Slopes for the fixed effects | $\beta_{i} \sim N(0.0, 3)$ | SM2b | SM2b |
