## Supplementary Material 3 for "Bayesian areal disaggregation regression to predict wildlife distribution and relative density with low-resolution data"

Supplementary Material 3 A1: 2018 Red deer posterior slopes for covariate fixed effects.

| **Parameter** | **Posterior mean** | **Posterior standard deviation** |
| --- | --- | --- |
| Artificial_surfaces | -1.90 | 1.21 |
| Broad_Leaf | 0.12 | 1.53 |
| Conifer | 0.04 | 1.41 |
| InlandWater | -0.03 | 1.41 |
| MixedForest | -0.05 | 1.49 |
| Moors_Heathland | -0.64 | 1.38 |
| NatGrassland | 0.38 | 1.62 |
| OpenSpaces | -0.18 | 1.64 |
| Pasture | 0.53 | 1.04 |
| PeatBogs | 4.01 | 1.52 |
| PlantedVegetation | -2.15 | 1.26 |
| Scrub | 0.64 | 1.59 |

Supplementary Material 3 A2: 2012 Red deer posterior slopes for covariate fixed effects.

| **Parameter** | **Posterior mean** | **Posterior standard deviation** |
| --- | --- | --- |
| Artificial_surfaces | -1.90 | 1.21 |
| Broad_Leaf | 0.12 | 1.53 |
| Conifer | 0.04 | 1.41 |
| InlandWater | -0.03 | 1.41 |
| MixedForest | -0.05 | 1.49 |
| Moors_Heathland | -0.64 | 1.38 |
| NatGrassland | 0.38 | 1.62 |
| OpenSpaces | -0.18 | 1.64 |
| Pasture | 0.53 | 1.04 |
| PeatBogs | 4.01 | 1.52 |
| PlantedVegetation | -2.15 | 1.26 |
| Scrub | 0.64 | 1.59 |

Supplementary Material 3 A3: 2006 Red deer posterior slopes for covariate fixed effects.

| **Parameter** | **Posterior mean** | **Posterior standard deviation** |
| --- | --- | --- |
| Artificial_surfaces | -1.9 | 1.2 |
| Broad_Leaf | -0.01 | 1.51 |
| Conifer | -0.24 | 1.37 |
| InlandWater | -0.31 | 1.4 |
| MixedForest | -0.06 | 1.49 |
| Moors_Heathland | 0.99 | 1.71 |
| NatGrassland | 1.09 | 1.79 |
| OpenSpaces | 1.17 | 2.06 |
| Pasture | -1.77 | 0.85 |
| PeatBogs | 2.67 | 1.02 |
| PlantedVegetation | -0.7 | 1.25 |
| Scrub | 1.13 | 1.53 |

Supplementary Material 3 A4: 2000 Red deer posterior slopes for covariate fixed effects.

| **Parameter** | **Posterior mean** | **Posterior standard deviation** |
| --- | --- | --- |
| Artificial_surfaces | -1.37 | 1.22 |
| Broad_Leaf | 0.07 | 1.52 |
| Conifer | 0.08 | 1.42 |
| InlandWater | -0.32 | 1.43 |
| MixedForest | 0.06 | 1.51 |
| Moors_Heathland | 0.26 | 1.58 |
| NatGrassland | 2.75 | 1.66 |
| OpenSpaces | 0.42 | 1.55 |
| Pasture | -1.95 | 0.95 |
| PeatBogs | 2.21 | 0.98 |
| PlantedVegetation | -0.7 | 1.21 |
| Scrub | -0.08 | 1.5 |

Supplementary Material 3 B1: 2018 Sika deer posterior slopes for covariate fixed effects.

| **Parameter** | **Posterior mean** | **Posterior standard deviation** |
| --- | --- | --- |
| Artificial_surfaces | -0.41 | 1.56 |
| Broad_Leaf | 0.46 | 2.08 |
| Conifer | 6.02 | 2.31 |
| InlandWater | 2.10 | 2.73 |
| MixedForest | -0.12 | 2.00 |
| Moors_Heathland | 3.51 | 1.40 |
| NatGrassland | 1.63 | 2.27 |
| OpenSpaces | 0.68 | 2.15 |
| Pasture | -2.46 | 0.82 |
| PeatBogs | -1.66 | 1.14 |
| PlantedVegetation | 1.31 | 0.82 |
| Scrub | -0.08 | 2.22 |

Supplementary Material 3 B2: 2012 Sika deer posterior slopes for covariate fixed effects.

| **Parameter** | **Posterior mean** | **Posterior standard deviation** |
| --- | --- | --- |
| Artificial_surfaces | -0.36 | 1.64 |
| Broad_Leaf | 0.59 | 2.05 |
| Conifer | 2.2 | 1.78 |
| InlandWater | 0.19 | 1.88 |
| MixedForest | 0.2 | 1.99 |
| Moors_Heathland | 4.07 | 1.48 |
| NatGrassland | 1.23 | 2.1 |
| OpenSpaces | 1.97 | 2.29 |
| Pasture | -2.81 | 0.89 |
| PeatBogs | -0.96 | 1.77 |
| PlantedVegetation | -1.42 | 1.33 |
| Scrub | 0.27 | 1.91 |

Supplementary Material 3 B3: 2006 Sika deer posterior slopes for covariate fixed effects.

| **Parameter** | **Posterior mean** | **Posterior standard deviation** |
| --- | --- | --- |
| Artificial_surfaces | -0.44 | 1.8 |
| Broad_Leaf | 0.45 | 2.03 |
| Conifer | 2.63 | 1.61 |
| InlandWater | -0.21 | 1.99 |
| MixedForest | -0.11 | 1.91 |
| Moors_Heathland | 1.74 | 1.99 |
| NatGrassland | -0.7 | 1.89 |
| OpenSpaces | 0.79 | 2.31 |
| Pasture | -2.61 | 0.96 |
| PeatBogs | 2.43 | 1.2 |
| PlantedVegetation | -1 | 1.22 |
| Scrub | 0.3 | 1.76 |

Supplementary Material 3 B4: 2000 Sika deer posterior slopes for covariate fixed effects.

| **Parameter** | **Posterior mean** | **Posterior standard deviation** |
| --- | --- | --- |
| Artificial_surfaces | -0.78 | 1.93 |
| Broad_Leaf | 0.27 | 2.03 |
| Conifer | -0.09 | 1.8 |
| InlandWater | -0.64 | 1.88 |
| MixedForest | 0.14 | 1.99 |
| Moors_Heathland | -0.8 | 1.86 |
| NatGrassland | 0.91 | 2 |
| OpenSpaces | 1.11 | 2.28 |
| Pasture | -2.3 | 1.12 |
| PeatBogs | 3.29 | 1.16 |
| PlantedVegetation | -1.6 | 1.41 |
| Scrub | 1.5 | 2.07 |

Supplementary Material 3 C1: 2018 Fallow deer posterior slopes for covariate fixed effects.

| **Parameter** | **Posterior mean** | **Posterior standard deviation** |
| --- | --- | --- |
| Artificial_surfaces | 1.62 | 1.36 |
| Broad_Leaf | 0.65 | 2.99 |
| Conifer | 7.58 | 2.15 |
| InlandWater | 0.00 | 3.35 |
| MixedForest | 1.49 | 3.05 |
| Moors_Heathland | 0.58 | 2.59 |
| NatGrassland | -0.34 | 2.91 |
| OpenSpaces | -0.93 | 2.78 |
| Pasture | 1.06 | 1.03 |
| PeatBogs | -2.03 | 1.78 |
| PlantedVegetation | -5.32 | 2.63 |
| Scrub | 6.17 | 2.30 |

Supplementary Material 3 C2: 2012 Fallow deer posterior slopes for covariate fixed effects.

| **Parameter** | **Posterior mean** | **Posterior standard deviation** |
| --- | --- | --- |
| Artificial_surfaces | -1.66 | 1.02 |
| Broad_Leaf | -0.18 | 2.37 |
| Conifer | 0.49 | 2.98 |
| InlandWater | 9.39 | 2.32 |
| MixedForest | 0.36 | 3.85 |
| Moors_Heathland | 0.66 | 3.02 |
| NatGrassland | 1.31 | 2.54 |
| OpenSpaces | -0.06 | 2.97 |
| Pasture | -0.73 | 2.93 |
| PeatBogs | 1.42 | 1.15 |
| PlantedVegetation | -2.49 | 2 |
| Scrub | -6.92 | 2.54 |

Supplementary Material 3 C3: 2006 Fallow deer posterior slopes for covariate fixed effects.

| **Parameter** | **Posterior mean** | **Posterior standard deviation** |
| --- | --- | --- |
| Artificial_surfaces | -1.8 | 1.09 |
| Broad_Leaf | 1.19 | 1.83 |
| Conifer | 0.83 | 3.05 |
| InlandWater | 7.04 | 1.82 |
| MixedForest | 0.89 | 4.31 |
| Moors_Heathland | 2.92 | 3.85 |
| NatGrassland | -0.45 | 2.95 |
| OpenSpaces | -1.41 | 2.77 |
| Pasture | -1.13 | 2.78 |
| PeatBogs | 0.77 | 1.29 |
| PlantedVegetation | -0.14 | 1.98 |
| Scrub | -8.27 | 2.35 |

Supplementary Material 3 C4: 2000 Fallow deer posterior slopes for covariate fixed effects.

| **Parameter** | **Posterior mean** | **Posterior standard deviation** |
| --- | --- | --- |
| Artificial_surfaces | -2.89 | 2.4 |
| Broad_Leaf | -0.03 | 2.99 |
| Conifer | 8.5 | 1.93 |
| InlandWater | 0.04 | 3.13 |
| MixedForest | 0.51 | 3.04 |
| Moors_Heathland | -1.46 | 2.61 |
| NatGrassland | -0.27 | 2.83 |
| OpenSpaces | -0.24 | 2.95 |
| Pasture | 2.28 | 1.2 |
| PeatBogs | -0.49 | 2.03 |
| PlantedVegetation | -6.14 | 2.17 |
| Scrub | 4.63 | 2.92 |
