## Supplementary Material Legends for "Bayesian areal disaggregation regression to predict wildlife distribution and relative density with low-resolution data"

Supplementary Material 1: Aggregated raw hunting bag return data for Irelands 3 deer species for the periods 2000, 2006, 2012 and 2018. Values indicate the number of red deer (top panel), sika deer (middle panel) and fallow deer (bottom panel) shot in a given year per county.

Supplementary Material 2: Priors' specification and posterior distribution estimated for all the parameters of the three species-specific (red, sika and fallow deer) models for 2000 (A), 2006 (B) and 2012 (C).. Spatial fields are defined in our model by their range (ρ) and standard deviation (σ). Priors are set on the range and standard deviation as penalised complexity (PC) priors and so the lower limit for the range and the upper limit for the SD are specified, together with a probability value for each. The fixed effects are defined by their slopes ($\beta_{i})$, and to these we set priors as a Gaussian distribution defined by its mean and SD. The posterior distributions of all parameters were obtained from the model and are described here by their mean and standard deviation.

Supplementary Material 3: Posterior fixed effect slopes of model covariates for 2000-2018 models for red deer (A1-A4), sika deer (B1-B4) and fallow deer (C1-C4).

Supplementary Material 4: Standard deviation of mean predictions for each of the 12 bayesian areal disaggregation models. Values indicate the lower 95% confidence intervals (A) and the upper 95% confidence intervals (B). Red deer are shown on the top panel, sika deer on the middle panel and fallow deer on the bottom panel.

Supplementary Material 5: Mean model predictions for the period 2000, 2006, 2012 and 2018 for red deer. Values indicate the relative density of red deer at 5x5km^2^ and are plotted on the logarithmic scale.

Supplementary Material 6: Mean model predictions for the period 2000, 2006, 2012 and 2018 for sika deer. Values indicate the relative density of sika deer at 5x5km^2^ and are plotted on the logarithmic scale.

Supplementary Material 7: Mean model predictions for the period 2000, 2006, 2012 and 2018 for fallow deer. Values indicate the relative density of fallow deer at 5x5km^2^ and are plotted on the logarithmic scale.
